# Autonomous MTOR activation in COPD airway epithelium impairs resolution through persistent mucus cell metaplasia and lysosome dysfunction

**DOI:** 10.64898/2026.09.15.751828

**Authors:** K. Kudrna, L.F. Vilches, X. Wang, J. Wang, T. Hugo, K.L. Bailey, J.D. Dickinson

**Affiliations:** Division of Pulmonary, Critical Care & Sleep Medicine, Department of Internal Medicine, College of Medicine, University of Nebraska Medical Center, Omaha, NE, USA; VA Nebraska-Western Iowa Health Care System, Research Service, Omaha, NE, USA

**Keywords:** Autophagy, COPD, IL-13, Lysosome, MTOR

## Abstract

**Background:** COPD is associated with persistent airway epithelial mucous cell metaplasia (MCM), mucin hypersecretion, and airway obstruction. Cytokines, such as IL-13, are well-known inflammatory drivers of airway MCM development. However, less is known about the factors that contribute to the impaired resolution of MCM in COPD. We hypothesized that MTOR activation in COPD airway epithelial cells (AECs) impairs autophagy and contributes to the delayed resolution of MCM.

**Methods:** AEC from COPD and non-diseased donors were grown under air-liquid interface (ALI) conditions and treated with IL-13 to promote MCM. Airway sections from COPD lung explants or non-diseased lung donors were utilized for immunohistochemical and immunostaining. Lysosomes were assessed using molecular probes and immunostaining.

**Results:** There was persistent MTOR-dependent MUC5AC immunostaining and epithelial hypertrophy in COPD AEC, which did not resolve after IL-13 withdrawal. MTOR substrate phosphorylation of RBS6, S6K1, and ULK1 was significantly elevated under baseline conditions, and at multiple timepoints independent of IL-13. Pharmacologic activation of MTOR led to increased IL-13-mediated MUC5AC levels in normal AEC but less robustly in COPD derived AEC. We hypothesized that persistent MTOR activation would reduce lysosome function and abundance in disease. COPD airways had reduced lysosome markers, LAMP1 and LAMP2 and fewer functional lysosomes by live-cell reporter probes. This lysosome deficiency in COPD AEC was partially rescued with MTOR inhibition.

**Conclusions:** We provide evidence to support an axis of autonomous MTOR activation and impaired lysosome function in the COPD airway epithelium. Persistent MTOR signaling is a molecular driver that contributes to persistent MCM.

**New & Noteworthy:** COPD is a leading cause of morbidity and mortality worldwide. Persistent mucous cell metaplasia (MCM), a key feature of this disease, is characterized by impaired resolution and prolonged exacerbations. IL-13 is increasingly recognized as a key mediator of viral- and environmental-driven MCM in COPD. Using IL-13-activated cultured COPD cells and airway sections from advanced COPD, we link autonomous MTOR signaling associated with impaired autophagy and lysosome function and persistent MCM.

## Introduction

The mammalian airway is lined by a layer of specialized epithelial cells that participate in mucociliary clearance. The plasticity of the airway epithelium has been well described during differentiation and in response to injury and inflammation^1^. In response to airway inflammation, basal cells and secretory cells further differentiate into large mucous-secreting cells that are overladen with mucin granules-morphologically termed goblet cells^2^. How airway goblet cells regulate size and morphology in the context of dynamic exogenous signaling is not completely understood. Muco-obstructive lung diseases (MOLD), including asthma, chronic obstructive pulmonary disease (COPD), and bronchiectasis, are the 3^rd^ leading cause of death in the U.S^3^. COPD is punctuated by exacerbations that lead to worsening of clinical symptoms and loss of lung function^4–7^. In response to acute and chronic inflammation from air pollution, cigarette smoke, allergens, and infections, patients with COPD develop epithelial changes, termed mucous cell metaplasia (MCM). The increase in MUC5AC during MCM contributes to mucus hyper-secretion and occlusion of small airways^8–11^. It is not surprising then that MUC5AC has been identified as a biomarker and an independent risk factor for disease progression in COPD^12^. Interleukin-13 (IL-13) is a type 2 cytokine that is a key driver of MCM in COPD^13^. IL-13 signaling via the IL-13/IL4^14^ receptor heterodimer leads to Stat6-dependent phosphorylation and increased MUC5AC production. In COPD patients, impaired resolution resides in aberrant airway basal cell differentiation^15^ and inflammation, which leads to persistent MCM, prolonging exacerbations and leading to long-term remodeling of the airway^15, 16^ ^6^. However, the mechanism for impaired MCM resolution is not yet understood.

Mammalian target of rapamycin (MTOR) is a nutrient-sensing signaling protein that activates cell growth, metabolism, and proliferation, and inhibits autophagy^17, 18^. The MTOR pathway is a highly conserved kinase signaling system that sits at the hub of this network. MTOR activation through phosphorylation of Ribosomal protein S6 kinase B1 (RPS6K1B) and Ribosomal Protein S6 (RPS6) leads to increased protein synthesis and anabolic metabolism required for cell growth and proliferation. Predictably, these cues via MTOR activation inhibit autophagy through phosphorylation of Unc-51 Like Autophagy Activating Kinase 1 (ULK1) and Autophagy-related protein 13 (ATG13) ^19–21^ as there is no need to recycle or conserve proteins. Autophagy is utilized in bulk protein breakdown in response to nutrient demands for new amino acids^22^. We recently identified that autophagy of mucin granules contributed to resolution of mucous metaplasia^23^ and that MTOR signaling regulates key features of transient IL-13-mediated MCM and resolution^24^ in airway epithelial cells. Here, we demonstrate that COPD-derived airway epithelial cells have impaired resolution from IL-13-mediated MCM due to persistent, elevated MTOR signaling. MTOR activation also impairs lysosome biogenesis in cultured COPD airway epithelial cells. These findings point to the critical role of MTOR activation in COPD pathogenesis by impairing resolution of epithelial MCM.

## Materials and Methods

### Human tissue and AEC culture

Histological airway cross sections of the conducting bronchi were obtained from COPD patients who underwent lung transplantation for end-stage disease. This protocol was approved by the UNMC IRB (#0077-25-EP). Normal airway sections were from deceased donors who were not selected for transplantation. The University of Nebraska Medical Center (UNMC) Division of Pulmonary, Critical Care, and Sleep Medicine maintains a tissue and cell bank from de-identified, non-diseased lungs obtained from Live On Nebraska, the local organ procurement agency. This protocol is approved by the UNMC IRB (#318-09-NH) and the Live On Nebraska ethics committee. Additional COPD airway cross-sections were obtained from Dr. Derek Byers, MD, Washington University School of Medicine (St. Louis, MO) under a protocol approved by the local institutional review board.

Normal human and COPD-derived airway epithelial cells (AEC) were derived from non-diseased human airways not suitable for transplant or COPD lung explant airways following clinically indicated lung transplantation as previously described^25, 26^. After enzymatic digestion of the airways, basal stem cells were expanded using BEGM media (Lonza 3170). Basal cells were then seeded on Corning transwell supports (Millipore Sigma CLS3401-48EA) for air-liquid interface (ALI) differentiation as previously described^25, 27^. AEC were differentiated using PneumaCult ALI media (Stem Cell 05001). After 21 days, healthy or COPD-derived AEC (hAEC or COPD AEC) were treated with human recombinant IL-13 (10ng/ml) in the basolateral chamber of the transwell. In order to activate MTOR signaling, MHY1485 was dissolved in DMSO and diluted to final concentration of 2.5 μM in normal AEC and COPD AEC. A subset of COPD AEC was treated with rapamycin 50nM for the indicated times in the figure legends.

### Immunofluorescence staining

Human AECs on Corning transwell membrane supports were harvested at the indicated time points. Basal media was removed, and 200 µL of 37-40°C 1% low melting point agarose was added to the apical surface. After cooling, the cells on membrane supports were fixed in 10% formalin and embedded in paraffin. AEC inserts and human airway sections were processed for immunostaining as previously described^23^. Antibodies for immunostaining included: mouse anti-MUC5AC 45M1 (Millipore Sigma MS-145-P1), rabbit anti-MUC5B (Atlas Antibodies HPA008246), rabbit anti-LAMP1 (Cell Signaling #D2D11), rabbit anti-LAMP2 (Cell Signaling 81197S), and rabbit anti-ACTIN (Cell Signaling 4970S). Image analysis was performed using a common threshold with ImageJ as previously described^23, 26, 27^.

### RPS6 immunohistochemistry

Slides were sectioned from formalin-fixed and paraffin-embedded tissue from healthy and COPD airway cross sections. They were incubated in 3 washes of xylene followed by 3 washes of isopropanol. Slides were then washed in deionized water, followed by antigen retrieval with Trilogy (Millipore-Sigma 922P-06). Slides were then rinsed in tris-buffered saline plus 0.1% triton X-100 (TBST), followed by endogenous peroxidases quenching with 3% hydrogen peroxide. Slides were then rinsed in tris-buffered saline with 0.5% Triton X-100 (TBST) and blocked with 10% normal goat serum (Vector S-1000) in phosphate-buffered saline (PBS) for 30 minutes at 37°C in a humidity chamber. Slides were then incubated with primary antibody Phospho-RPS6 (Cell Signaling 2215S) diluted in 0.5% bovine serum albumin (BSA) in PBS overnight at 4°C. Slides were then rinsed in TBST, followed by biotinylated goat secondary antibody (Vector BA-1000) in the same blocking buffer. After washing the slides in PBS, ABC reagent (Vector PK-7100) was used, followed by chromogen substrate staining-DAB (Vector SK-4105). Slides were washed in PBS and counterstained with hematoxylin, washed in water, and dehydrated with increasing concentrations of ethanol incubations and then xylene.

### Immunoblotting

MTOR substrate proteins were measured from normal human AEC and COPD AECs. NP-40 buffer supplemented with protease/phosphatase inhibitors was used for lysis to preserve phosphorylation sites as previously described^24^. The following antibodies were used for detection of protein on PVDF membranes following transfer: total MTOR (Cell Signaling 2972S), P70S6K total and T389 phosphorylation (Cell Signaling 9202S and 9205S), Ribosomal S6 total and S240, 244 phosphorylation (Cell Signaling 2217S and 2215S), ULK1 total and S757 phosphorylation (Cell Signaling 8054S and 14202S) and LC3 was detected by LC3B (Sigma-Aldrich L7543). Signal was quantified with Infrared (IR) labeled secondary antibodies (LiCor) and normalized to total cytoplasmic protein. In order to measure 4 different MTOR substrate proteins from the same Western blot, the membrane was cut prior to the blocking step, based on molecular weight markers.

Detection of MUC5AC by immunoblotting was performed from normal and COPD AEC cell lysates. Cells were lysed in a cocktail of phosphate-buffered saline (PBS) with protease inhibitors and then sonicated for 5 seconds until lysate was clear. Lysates were then centrifuged at 15,000 g for 30 minutes as previously described^23, 24^. After blocking in a solution of 5% milk in PBS, membranes were probed with mouse MUC5AC - 45M1 (Millipore Sigma MS-145-P1) antibody overnight at 4°C on a rocker. After PBS washes, the membranes were probed with goat anti-mouse IR conjugated antibody (LiCor), and bands were normalized to total protein stain for quantification.

### Live cell lysosome quantification

Healthy and COPD-derived basal cells were cultured in BEGM media until confluent. A subset of COPD cells was treated with rapamycin 50nM or DMSO vehicle for 24 hours. A second subset was treated with 5% cigarette smoke extract (CSE) for 24 hours, prepared as previously described^28^. Prior to Lysotracker probe loading, cells were washed with sterile Hanks buffered saline solution (HBSS). Then 1 μM of diluted LysoTracker™ Red DND-99 (ThermoFisher L7528) in warmed sterile HBSS was added along with Hoechst 33342 nucleic acid stain (ThermoFisher H3570). AEC were cultured for 40 minutes at constant 37°C and 5% CO2. AEC were then washed in warm sterile HBSS prior to imaging.

### Statistical Analysis

Each statistical test is described in the corresponding figure legend. T-test or ANOVA with Tukey’s pairwise comparison was used for normally distributed data. Mann-Whitney U test was used to compare ranks in non-normally distributed data. P<0.05 was considered significant. All comparisons were unpaired analyses.

## Results

### Persistent MTOR-dependent MUC5AC levels following IL-13-mediated mucous cell metaplasia in COPD airway epithelial cells

IL-13 is a significant inflammatory driver of airway MCM and contributes to both severe asthma and COPD.^13, 29^ It leads to airway obstruction by mucus obstruction and airway wall thickening^30, 31^. The factors that thwart resolution of MCM in COPD are not yet well described. We recently reported that in normal airway epithelial cells (AEC), IL-13-mediated MCM, cellular hypertrophy, and hyperplasia largely resolved 7 days after withdrawal of the cytokine^24^. We, therefore, next examined the impact of this model in AEC derived from end-stage COPD airways. These COPD AEC differentiate under air-liquid interface conditions and produce abundant basal, ciliated, and secretory cells. Unlike normal AEC, we found that there was persistent airway epithelial hypertrophy and elevated MUC5AC levels in COPD AEC even after IL-13 withdrawal (**Fig. 1A-C**). Thus, COPD AEC demonstrate impaired IL-13-mediated MCM resolution. We previously demonstrated that MTOR activity was dynamically associated with IL-13-mediated MCM and resolution in normal AEC^24^. Therefore, we hypothesized that failure of resolution was related to persistent MTOR signaling. Pharmacologic inhibition of MTOR with rapamycin led to normalization of airway epithelial hypertrophy and MUC5AC levels during resolution (**Fig. 1B-E**). These findings indicate that COPD AEC have dysregulated resolution of IL-13-mediated MCM related to persistent MTOR activation.

**Figure 1:**
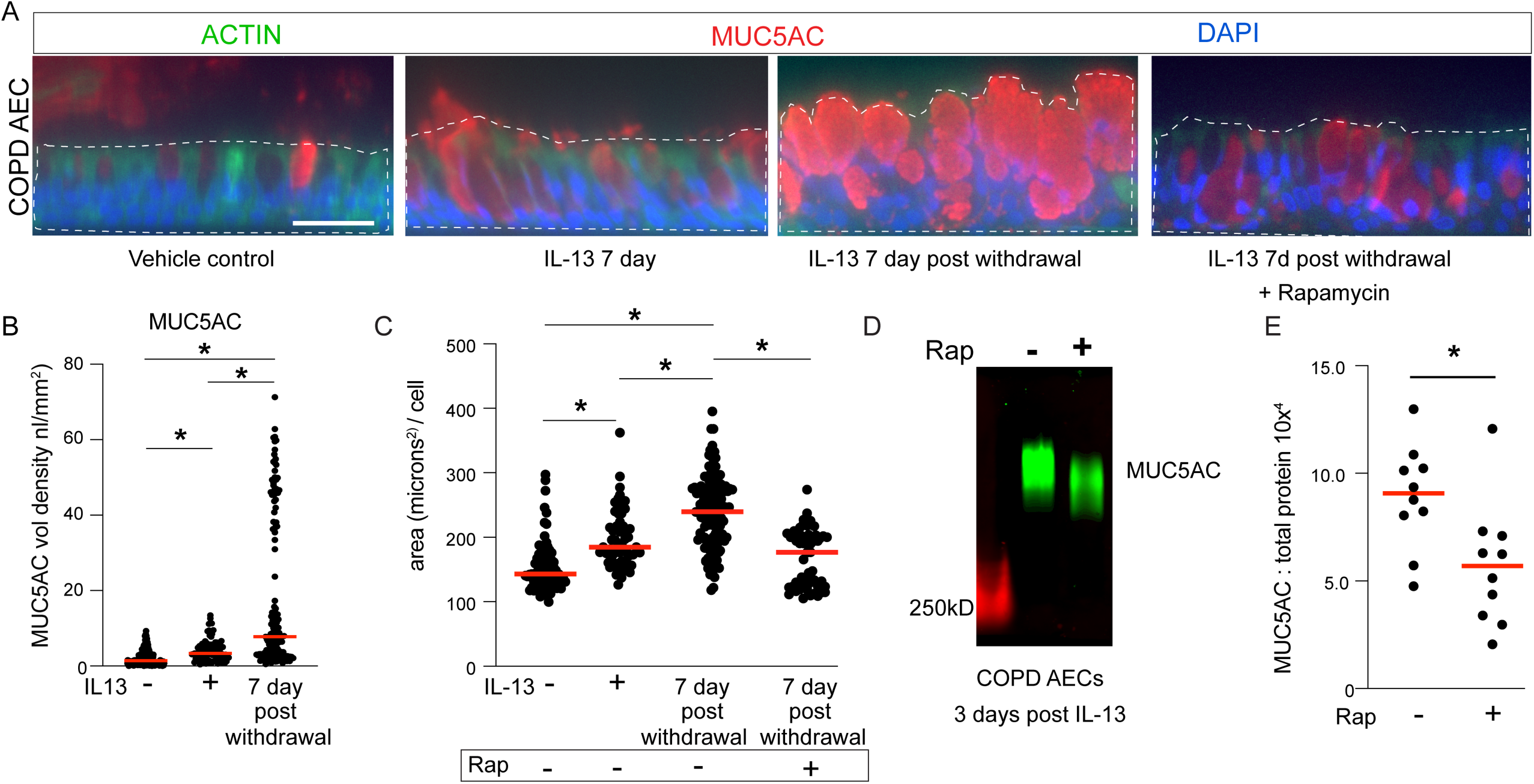
Persistent MTOR-dependent IL-13-mediated mucous cell metaplasia (MCM) in COPD airway epithelial cells. **A**) Representative immunostaining for MUC5AC and cytoplasmic ACTIN under homeostatic conditions, after 7 days of IL13, during resolution 7 days after withdrawal of IL-13, and after treatment with Rapamycin 50nM for 48 from day 5 to day 7 post IL-13 withdrawal. DAPI for nuclear counterstain. Dashed lines depict measured cytoplasmic area per image. Scale bar represents 50 microns. **B,C**) Quantification of cytoplasmic area as outlined by ACTIN staining per cell and MUC5AC / volume density per airway segment for COPD AEC. Red bar is median area, n=10-12 images per sample from 5 unique donors. **D,E**) Representative MUC5AC immunoblot from COPD AEC cell lysates 3 days after IL-13 withdrawal treated with vehicle or Rapamycin 50nM for 48 hours with corresponding quantification. Red bar is median MUC5AC volume density, n=10 from 2 unique COPD donors. * P<0.05 by ANOVA with Tukey’s Multiple Comparison test for pairwise comparison (**B,C**) or unpaired T-test (**E**).

### Autonomous MTOR signaling is dysregulated in COPD airway epithelium

MTOR is a highly conserved signaling hub that integrates exogenous and endogenous cellular signaling induced by infections, metabolic changes, and inflammatory cues^32^. We previously showed that IL-13 transiently activates MTOR substrate phosphorylation at RPS6, RPS6KB1, and ULK1. Withdrawal of IL-13 led to a corresponding decrease in phosphorylation of these proteins in normal AEC^24^. We hypothesized that the lack of observed MCM resolution in COPD AEC was driven by dysregulated MTOR signaling. We cultured normal AEC and COPD AEC under ALI conditions until fully differentiated (21-28 days). We found a significant increase in phosphorylated MTOR substrates (RPS6, RPS6KB1, and ULK1) under baseline ALI conditions (**Fig. 2A,B**). At multiple timepoints with IL-13 activation and even 7 days after IL-13 withdrawal, we found significantly increased phosphorylation of RPS6, S6K1, and ULK1 in COPD AEC compared to normal AEC (**Fig. 2B-E**). In normal airway AEC, MTOR substrate phosphorylation was dependent on IL-13 activation^24^. Despite having elevated MTOR substrate phosphorylation at each time point, COPD AEC did not show the same IL-13-dependent increase. This suggests an epithelial autonomous MTOR activation in COPD that augments existing IL-13-mediated MCM.

**Figure 2:**
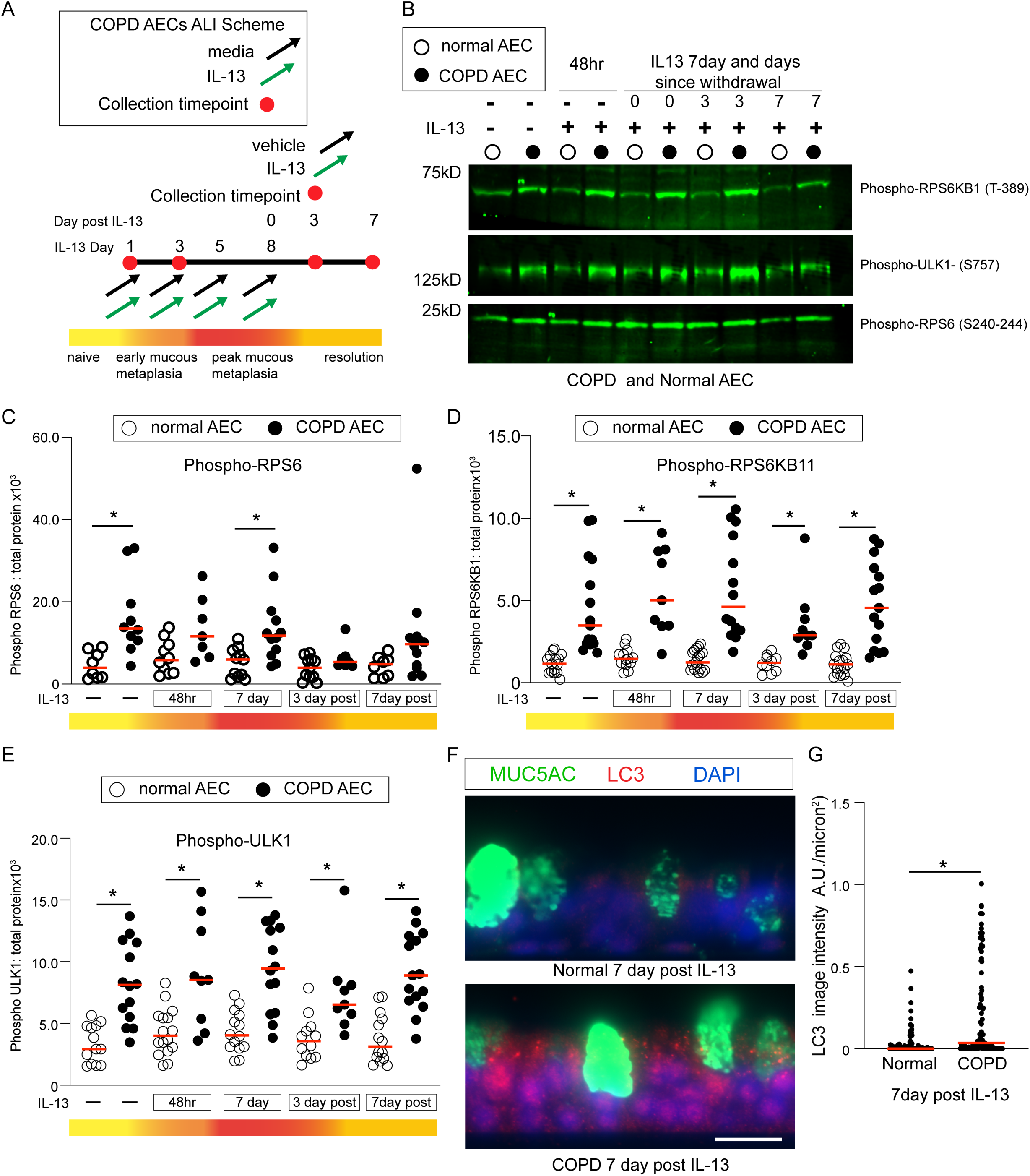
Autonomous MTOR activation in COPD AEC independent of IL-13. **A**) Schematic showing model of IL-13-mediated MCM and resolution after cytokine withdrawal for normal and COPD-derived AEC. **B**) Representative immunoblot for phosphorylated MTOR substrates RPS6KB1, RPS6, and ULK1 from normal and COPD-derived AEC at the indicated timepoints of IL-13-mediated MCM and IL-13 withdrawal. **C,D,E**) Quantification of phosphorylated RPS6, RPS6KB1, and ULK1 normalized to total protein. N=8-13 from 8 unique normal or COPD donors. * p<0.05 for Kruskal-Wallis test with Dunn’s multiple comparison tests. **F**) Representative immunostaining for autophagy marker, LC3, from normal and COPD-derived AEC. **G**) Quantification of LC3 puncta intensity per airway section area (microns^2^). N=103 airway images from 5 unique normal donors and 108 airway images from 5 unique COPD donors. * p<0.05 for Mann-Whitney Test.

ULK1 phosphorylation at position serine 757 is known to inhibit autophagy activation^20^. Our data suggested that dysregulated MTOR-phosphorylation of ULK1 impairs autophagy. Autophagy protein LC3 is a marker of mature autophagosome membranes. Increased LC3 staining denotes either increased autophagy and/or lysosome inhibition. During resolution, we found elevated LC3 immunostaining in differentiated COPD AEC compared to differentiated normal AEC (**Fig. 2F,G**). In conjunction with elevated phosphorylation of ULK1, these findings indicate that failure of resolution of MCM in COPD is associated with persistent elevation of MTOR signaling and impaired autophagy. To confirm the significance of persistent elevation of MTOR activity in cultured COPD AEC under ALI conditions, we examined phosphorylated MTOR substrate proteins, RPS6 in airway epithelial sections from COPD explant lungs and normal non-diseased donor lungs not suitable for lung transplant. There was a significant increase in phosphorylated-RPS6 in COPD airway epithelium compared to normal airways (**Fig. 3A,B**).

**Figure 3:**
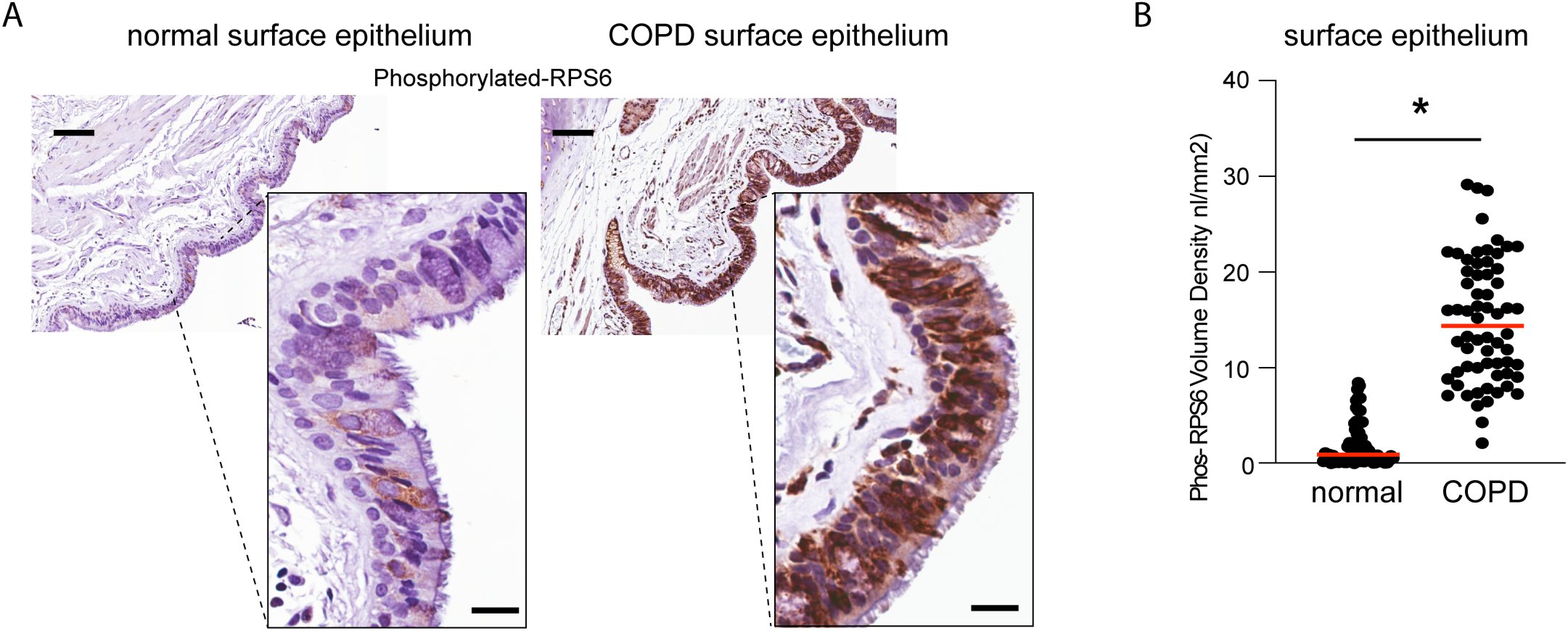
MTOR activity increased in COPD airways. **A**) Representative immunohistochemical staining for phosphorylated RPS6 from non-diseased and COPD airways sections. Scale bar is 20 microns. **B**) Quantification of Phos-RPS6 volume density from normal and COPD airways. N=10-15 images per donors, 5 donors per condition * p<0.05 by Student T test.

### MTOR activation and IL-13 activation lead to synergistic impact on MUC5AC mucous cell metaplasia

Based on the temporal association between persistent MUC5AC MCM and MTOR activation in COPD AEC following IL-13 activation and withdrawal, we examined the impact of pharmacologic MTOR activation under IL-13-dependent and independent conditions. MTOR activator MHY-1485 increases MTOR signaling^33^ and blocks lysosome-autophagosome fusion^34^. Therefore, we cultured normal AEC under ALI conditions until fully differentiated at day 21-28. Normal AEC were then treated with DMSO vehicle, IL-13 (10ng/mL), MHY-1485, or the combination of MHY1485 and IL-13 for 7 days (4 media changes) (**Fig. 4A**). The addition of MHY-1485 led to a significant increase in the phosphorylation of RPS6KB1 (**Fig. 4B,C**) but only in IL-13-activated AEC. We next examined the impact of transient pharmacological MTOR activation only during IL-13-mediated MCM in ALI-differentiated normal AEC. There was a significant synergistic impact with combined MTOR pharmacologic treatment and IL-13 activation. MUC5AC levels by immunostaining were significantly increased with IL-13 plus MHY-1485 compared to IL-13 alone (**Fig. 4D,E**). MHY-1485 alone had no impact on MUC5AC immunostaining levels. In order to confirm the synergistic impact of MTOR activation by MHY-1485 and IL-13 treatment, we utilized mucin immunoblotting of normal AEC lysates and found a comparable significant increase in MUC5AC levels from cell lysates compared to IL-13 (**Fig. 4F,G**). MTOR activation prolonged IL-13 mediated mucous cell metaplasia (**Fig. 1D,E).** We then sought to determine if induced MTOR activation would similarly augment MUC5AC MCM in COPD AEC differentiated under ALI conditions. Immunoblotting of COPD AEC lysates revealed a similar synergistic, increase in MUC5AC levels after IL-13 plus MHY-mediated MTOR activation (**Fig. 4H,I**). The magnitude of this increase was approximately a 2-fold change compared to a 4-fold change normal human AEC (**Fig. 4G**). Immunostaining of COPD derived AEC under ALI conditions, confirmed the increase in MUC5AC levels was due to cytoplasmic MUC5AC (**Fig. 4J**). These findings suggest that MTOR activation further increases the impact of IL-13 on airway mucous cell metaplasia, although there appears to be a ceiling to cellular response to induced pharmacologic or IL-13-mediated MTOR activation in COPD cells.

**Figure 4:**
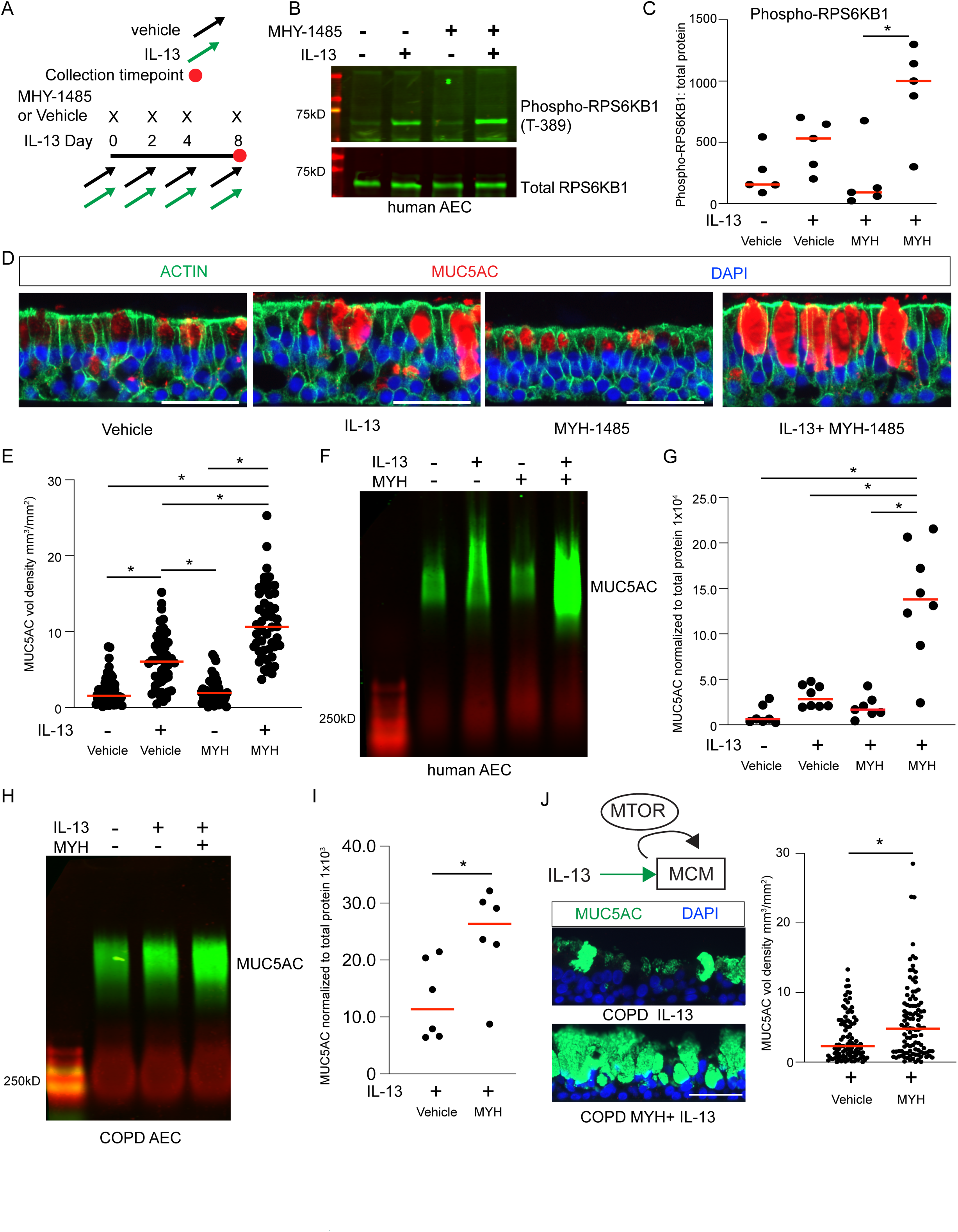
MTOR and IL-13 activation leads to synergistic increase in MUC5AC mucous cell metaplasia. **A**) Treatment scheme for IL-13 and MTOR pharmacologic activator, MYH-1485 in normal AEC under ALI conditions. **B**) Representative immunoblot for total and phosphorylated RPS6KB1 from human AEC in duplicate using conditions described in panel **A**. **C**) Quantification of Phosphorylated RPS6KB1 normalized to total protein. N=5 from 4 unique hAEC donors. **D**) Representative immunostaining of human AEC for MUC5AC and cytoplasmic ACTIN to mark cytoplasmic borders with DAPI for nuclear counterstain. Scale bar equals 50 microns. **E**) Quantification of MUC5AC volume density from 10-15 microscopic images per microscopic slide from N=5 unique normal AEC donors. * p<0.05 by ANOVA with Tukey’s multiple comparison test. **F,G**) Representative immunoblot and quantification for MUC5AC from 8 samples from 5 unique human AEC donors. MUC5AC normalized total protein from lysate. * p<0.05 by ANOVA with Tukey’s multiple comparison test. **H,I**) Representative immunoblot for MUC5AC from COPD-derived AEC under ALI conditions with vehicle, IL-13, and IL-13 plus MYH-1485. Quantification of MUC5AC levels normalized to total protein levels. N=6 from 4 unique COPD AEC donors treated with IL-13 or IL-13+MYH-1485. * p<0.05 by Mann Whitney test. **J**) Representative MUC5AC immunostaining with DAPI counterstain for COPD AEC treated with either IL-13 or IL-13+MYH-1485 for 7 days. Quantification of MUC5AC volume density from 97 and 107 microscopic images from 7 unique COPD donors. * p<0.05 by Student T test

### MTOR activity impairs COPD airway epithelium cells’ lysosomes

Lysosomes are key organelles that degrade cytoplasmic proteins and organelles delivered by the autophagosome. They also play key metabolic and inflammatory functions and are dysregulated in disease, including COPD^35^. Lysosome formation and function are regulated by MTOR. MTOR activation leads to impaired lysosome formation through phosphorylation and cytoplasmic sequestration of the transcription factor TFEB^36^. We have found elevated MTOR signaling in COPD AEC (**Fig. 2 and 3**). We therefore examined the abundance and function of lysosomes in the COPD airway epithelium. Cigarette smoke exposure is the predominant environmental insult that drives COPD pathogenesis. We cultured COPD AEC with and without 5% cigarette smoke extract (CSE) for 24 hours. CSE induced a significant decrease in functional lysosomes as measured using a live cell molecular probe, Lysotracker Red^TM^ (**Fig. 5A,B**). The associated decrease in functional lysosomes correlated with increased LC3 II levels from COPD AEC (**Fig. 5C,D**). This data suggests that cigarette smoke exposure impairs COPD lysosome function and impairs autophagy flux. We next sought to verify this lysosome impairment in COPD airways and cultured cells. Using the lysosomal membrane markers, LAMP1 and LAMP2, we found diminished LAMP1- and LAMP2-positive lysosomes in COPD airways relative to non-diseased airways (**Fig. 6A-D**). Interestingly, we found that LAMP2 staining was predominantly focused on the MUC5AC-positive goblet cells in both the normal and COPD airways (**Fig. 6C inset, E**). However, COPD MUC5AC-positive goblet cells had significantly reduced LAMP2 staining compared to normal MUC5AC-positive goblet cells. This suggests that LAMP2-labeled lysosomes play an important role in MUC5AC-positive secretory cells and are impaired in COPD goblet cells.

**Figure 5:**
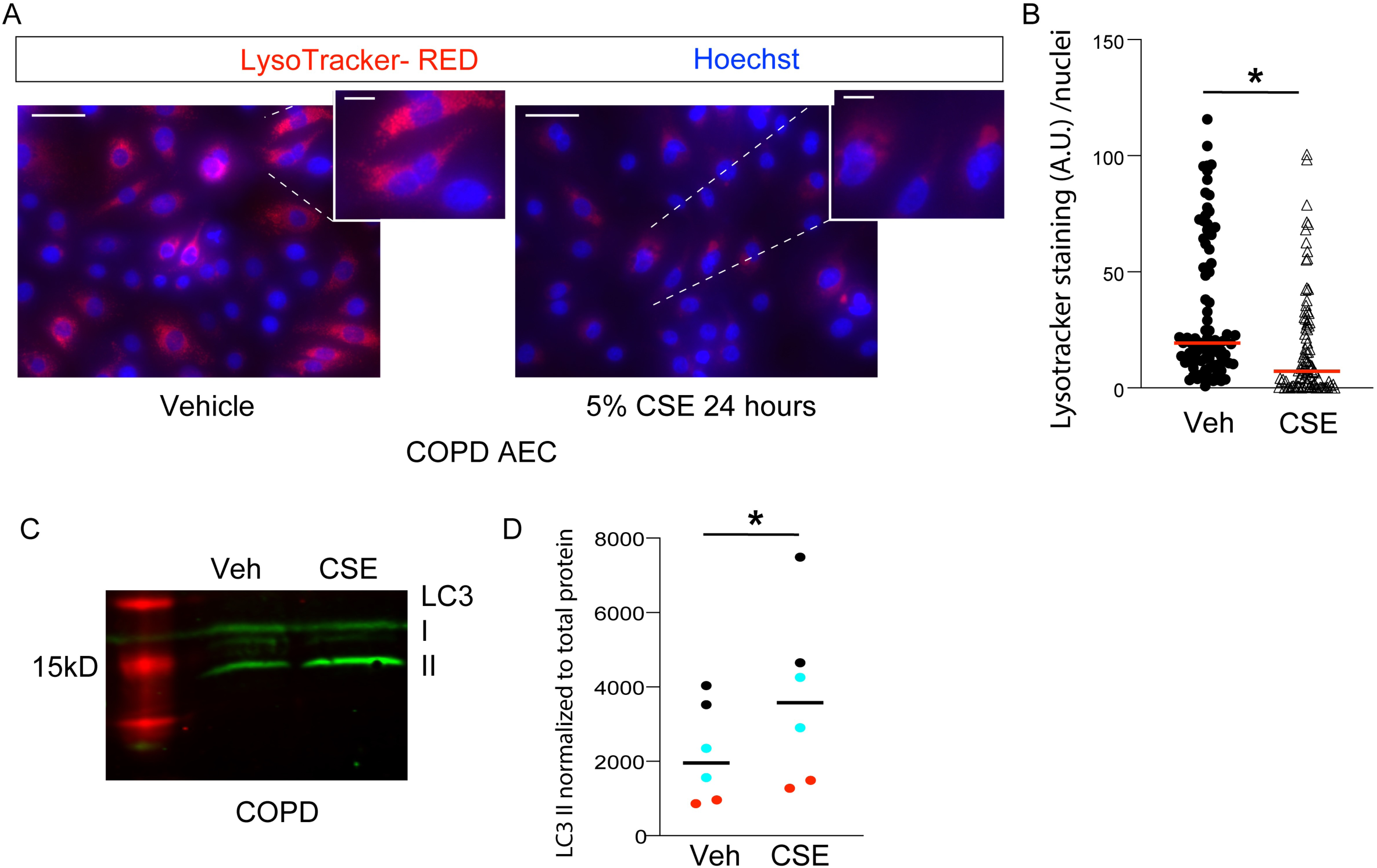
Cigarette smoke extract decreases functional lysosomes in COPD AEC. **A**) Representative images of COPD AEC basal cells either treated with vehicle or 5% Cigarette Smoke Exposure (CSE) for 24 hours. Scale bars are 50microns and 10 microns in inset. **B**) Quantification of Lysotracker RED staining per nuclei. N=121 and 122 images from 6 unique COPD donors. * P<0.05 by Mann-Whitney Test. **C**) Representative LC3 immunoblot for COPD AEC with and without CSE. **D**) Quantification of LC3 II normalized to total protein. N=6 from 3 unique COPD donors (each donor color coded). * P<0.05 by Paired T-test.

**Figure 6:**
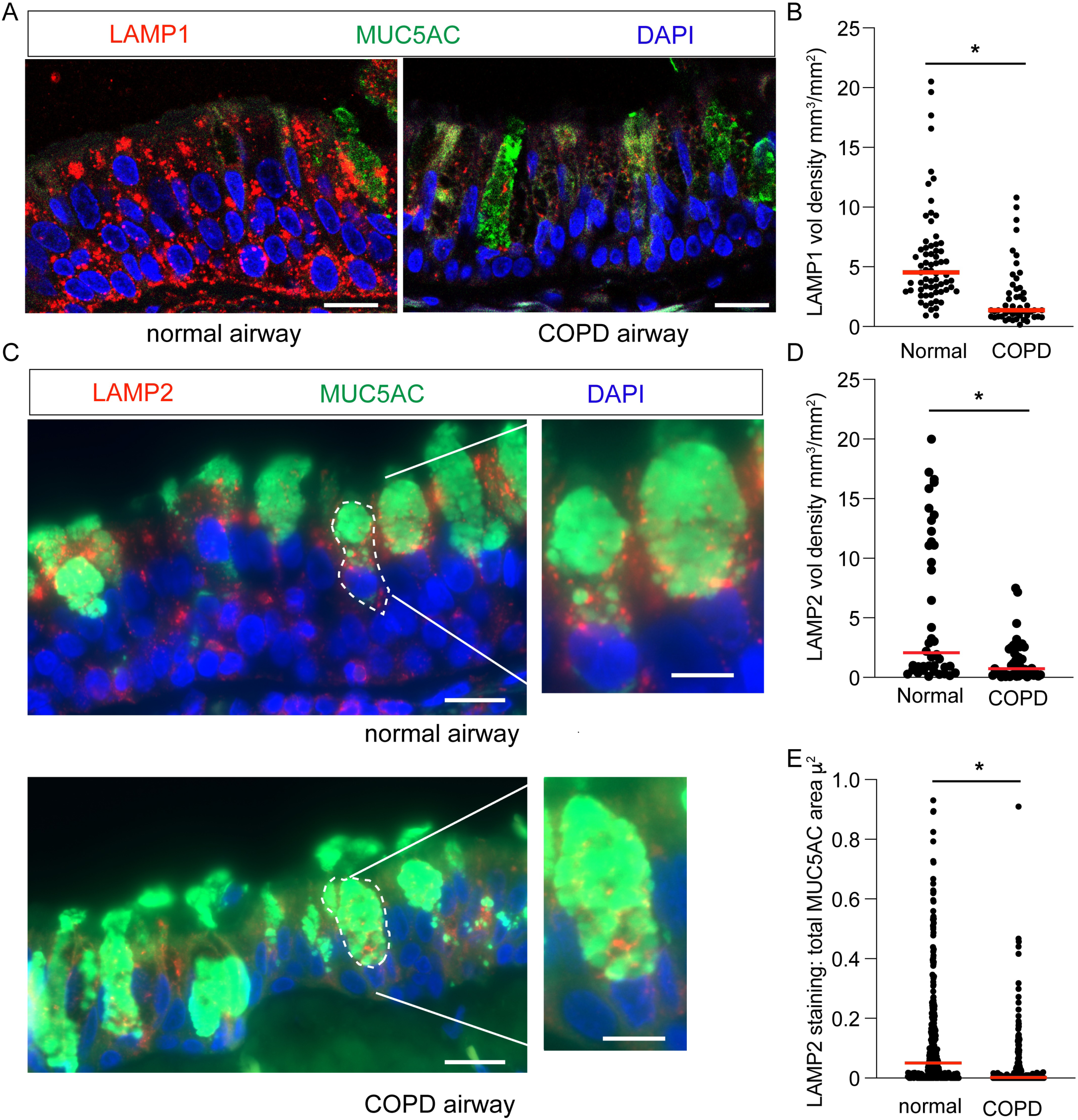
Reduced LAMP1 and LAMP2-labeled lysosomes in COPD airways. Representative image of lysosome markers, LAMP1 (**A**), LAMP2 (**C**), MUC5AC, and DAPI for nuclear counterstain in cross sections of non-COPD airway control and COPD airways. **B,D**) Quantification of LAMP1 and LAMP2 volume density per airway segment. For part A,B, N=67 and 50 microscopic images from 4 unique normal and COPD donors. For part C,D, N= 40 and 44 microscopic images from 3 normal and 4 unique COPD donors. Scale bar is 20 microns and 10 microns in inset image. **E**) Quantification of LAMP2 staining intensity (A.U.) per MUC5AC goblet cell area microns^2^ (denoted by dashed line). N=3 normal donors (361 unique goblet cells) and 4 COPD donors (274 unique goblet cells). * P<0.05 by Mann-Whitney Test for parts B,D,E.

We hypothesized that increased MTOR signaling may suppress lysosome abundance and function in COPD. We cultured normal and COPD basal cells to examine functional lysosomes. We found decreased Lysotracker Red^TM^ in COPD basal cells compared to normal basal cells (**Fig. 7A,B**). Brief treatment with the mTOR inhibitor rapamycin significantly increased the number of COPD basal cell functional lysosomes (**Fig. 7A,B**). Next, we examined COPD AEC differentiated under ALI conditions. Treatment with rapamycin for 48 hours also led to an increase in levels of lysosome marker LAMP2 (**Fig. 7C,D**) in COPD AEC. These data indicate that dysregulated MTOR signaling is central to impaired lysosome function in COPD airway epithelial cells.

**Figure 7:**
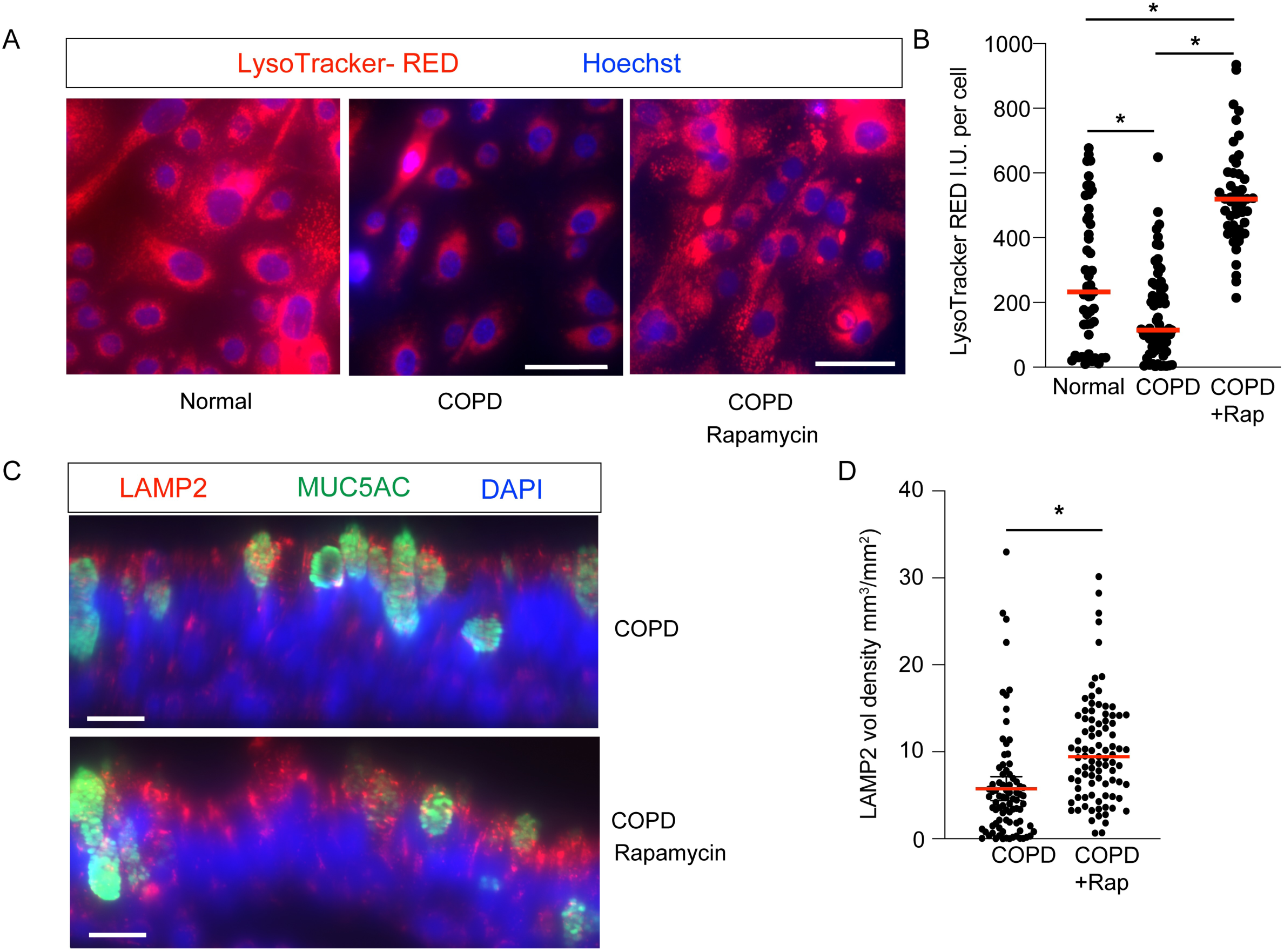
MTOR activation suppresses lysosomes in COPD airway epithelial cells. **A**) Representative images of LysoTracker-RED lysosome live imaging stain in normal and COPD basal AEC with Hoechst for nuclear counterstain. Scale bar 50 microns. N= 49 images from 3 normal donors and 69 images from 4 COPD donors. A subset of COPD AEC were treated with rapamycin 50nM for 24 hours. N=47 images from 4 COPD donors. **B**) Quantification of Lysotracker-RED staining per nuclei. **C**) Representative LAMP2, MUC5AC immunostaining from COPD AEC cultured under ALI conditions and treated with Vehicle or Rapamycin 50nM for 48 hours. **D**) Quantification of LAMP2 volume density. N=4 COPD donors from 83 and 91 microscopic images, respectively, for each condition. * P<0.05 by Mann-Whitney Test.

## Discussion

Mucous cell metaplasia (MCM) is a key pathologic feature of airway diseases including COPD. COPD airway epithelium demonstrates persistent MCM revealed by longitudinal clinical studies demonstrating airway wall thickening and mucus plugging on chest CT^37^. MCM is confirmed histologically by epithelial hypertrophy and goblet cell hypertrophy in COPD sections^8, 9^. Airway goblet cells in COPD have an increased number of stored cytoplasmic mucin granules. Persistent airway MCM leads to mucous hyper-secretion of these stored granules and the development of mucus plugging, which is now recognized as an independent predictor of outcomes in COPD^37^. In addition, MUC5AC sputum levels are an effective biomarker of disease burden^38^. Finally, IL-13 is now recognized as a key driver of persistent MCM in COPD^13^ and can be targeted by novel biologic agents^29^. Therefore, understanding the key molecular drivers of persistent MCM is important for disease-modifying therapies. Abnormal basal cell differentiation and function have long been recognized as a key feature in COPD^15, 39, 40^. How these basal cells respond to IL-13-mediated inflammation in relation to MCM and mucus plugging has not yet been fully explored. We demonstrate that, in contrast to normal AEC in which IL-13 increases MTOR activation, MTOR activation in COPD AEC appears to be autonomous and acts as a molecular driver that augments IL-13-mediated MCM. Interestingly, we found that either genetic^24^ or pharmacologic activation of MTOR significantly increased IL-13-mediated MCM and cytoplasmic stored MUC5AC granules in goblet cells. This response was less robust in COPD AEC. This may be related to the autonomous MTOR-mediated phosphorylation substrates at baseline. Therefore, MTOR signaling is fundamentally dysregulated in COPD airway epithelium and leads to persistent elevation of LC3 levels reflecting impaired autophagy and lysosome dysfunction.

We recently identified that deletion of autophagy gene Atg16L1 in mouse secretory cells blocked autophagy and led to a cytoplasmic accumulation of MUC5AC mucin granules during resolution of MCM. Further, we identified that mucin granules fuse with lysosomes in an autophagy-dependent fashion. Therefore, lysosome function is a key regulator of elimination of excess mucin granules following MCM. Here we demonstrate that COPD AECs have reduced functional lysosomes and reduced staining of lysosome markers both in undifferentiated basal cells and intact airway epithelium from end-stage COPD airways. Our data confirms and expands on a recent finding that LAMP2 levels were decreased in COPD epithelium at baseline and after CSE exposure^41^. We found that LAMP2 levels in COPD airways, particularly in MUC5AC positive cells. LAMP2 is an important regular of lysosome function and selective autophagy^42^. Interestingly, lysosome numbers and LAMP2 levels could be rescued by MTOR inhibition using rapamycin in both basal cells and differentiated COPD airway epithelial cells. This suggests that the balance of MTOR and autophagy in the airway epithelium may directly regulate LAMP2 levels on the lysosome. Whether LAMP2 is a specific receptor for selective autophagy of mucin granules in the airway epithelial secretory cells remains to be determined.

Treatment of COPD has primarily relied on symptom relief with bronchodilators and prevention of exacerbations with inhaled corticosteroids. Yet these therapies do not improve survival or modify disease progression. Smoking cessation is key for the improvement of symptoms and reduction of the rate of decline in lung function. Yet, even in ex-smokers, MCM persists along with exacerbations. Therefore, there is a need for disease-modifying therapies to nudge the reprogrammed airway epithelium toward resolution. In a subset of COPD patients with elevated peripheral blood eosinophils, dupilumab, which targets the IL-13/IL-4 receptor, reduced exacerbations and modestly increased lung function^29^. It is not yet known the impact of dupilumab on mucus plugging and airway wall thickness. Nor is it known the durability of response. Our data here implies that IL-13-mediated MCM of the airway persists after the cytokine is removed due to autonomous MTOR activation. Novel treatments will likely be required to restore normal mucociliary clearance and tissue resolution in COPD. MTOR signaling offers an attractive therapeutic target to mitigate the long-term impacts of MCM in COPD. However, recognized clinical pulmonary toxicity of MTOR inhibitors such as sirolimus^43^ may limit the therapeutic option of this pathway. More selective agents that drive autophagy-mediated degradation of mucin granules in COPD airway hyperplastic goblet cells are needed as potential disease-modifying therapies. These studies were performed using AEC derived from end-stage COPD lung disease. Additional studies are needed from AEC derived from various stages of COPD severity to determine the longitudinal impact of MTOR dysregulation in both active, prior, and non-smokers with COPD.

## Conclusion

COPD airway epithelium is characterized by impaired resolution of IL-13-mediated mucous cell metaplasia with MTOR-dependent prolonged MUC5AC goblet cell hyperplasia and epithelial hypertrophy. MTOR is dysregulated in COPD with persistent elevation of phosphorylated substrates independent of IL-13 activation. Finally, autonomous MTOR activation in COPD downregulates key markers of lysosome structure and function. Targeting MTOR activation may provide a new pathway for reshaping persistent MCM in COPD.

## Author Approvals

All authors have seen and approved the manuscript, and it hasn’t been accepted or published elsewhere.

## Data Availability

Source data for this study are openly available at https://doi.org/10.5281/zenodo.21476552.

---

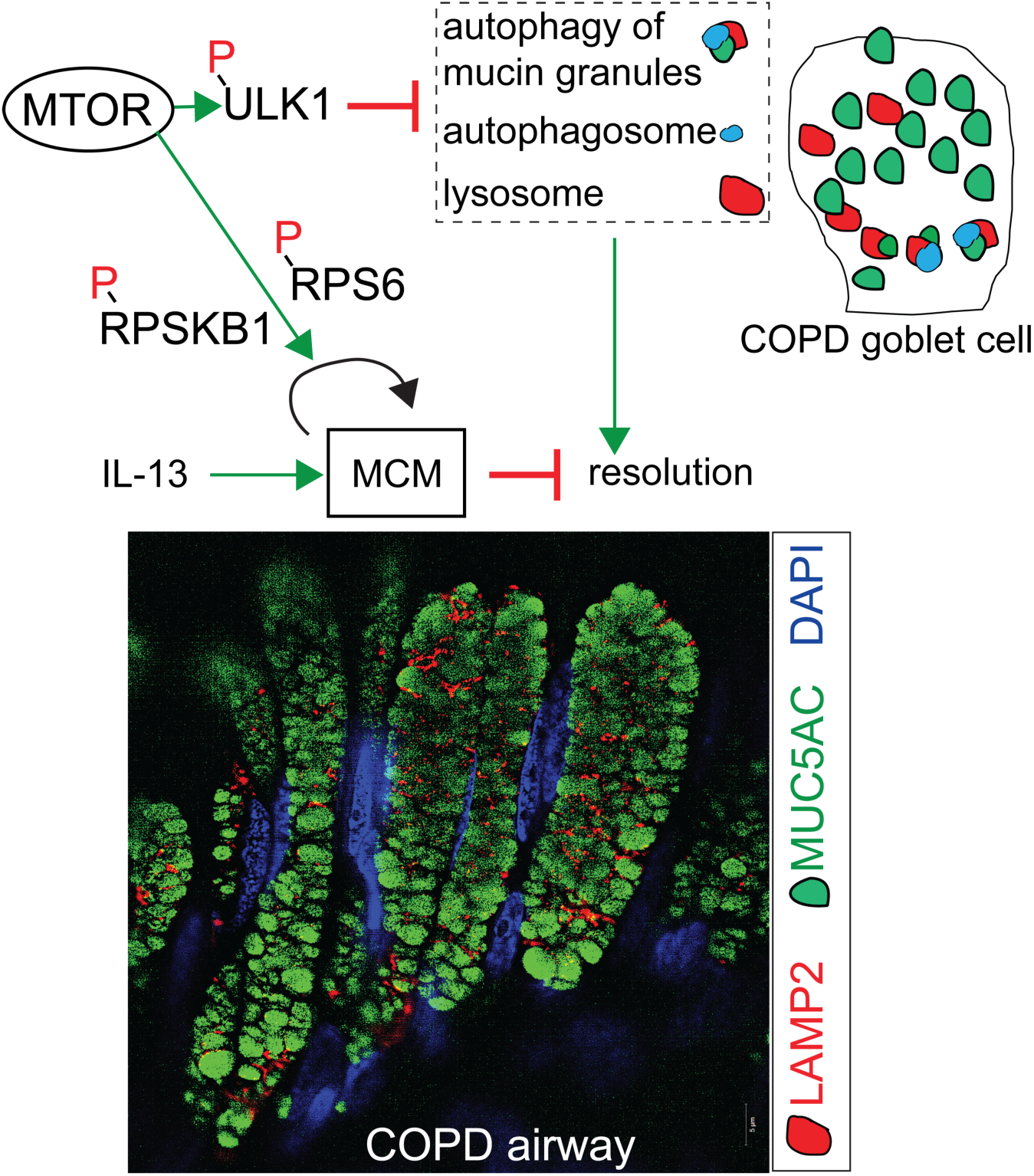

